# Source-sink dynamics can maintain mismatched range and bioclimatic limits even at large spatial scales

**DOI:** 10.1101/2020.12.05.413153

**Authors:** Nikunj Goel, Timothy H. Keitt

## Abstract

Bioclimatic models assume that at broad spatial scales, climate is the primary determinant of species distribution. Meanwhile, processes such as source-sink dynamics can be ignored because they are thought to manifest at length scales comparable to species mean dispersal distance. We present a reaction-diffusion model to show species can use sink patches near the bioclimatic (or niche) limit as stepping-stones to occupy sinks much further than the mean dispersal distance, thereby extending the distribution far beyond the bioclimatic envelope. This mismatch between geographical and bioclimatic limits is mediated by the shape of the bioclimatic limit and may be significant for low growth sensitivity and fast dispersal life strategy. These findings challenge one of the core assumptions of the bioclimatic models. Therefore, we advocate that biogeographers consider the role of dispersal when using bioclimatic models to generate inferences about the ecological and evolutionary processes that determine the distribution of biota.

## Introduction

All species are geographically limited. Understanding the mechanism that limits species distributions is a central challenge in biogeography theory (Gaston 2003, Holt and Keitt 2005). Traditionally, biogeographers have argued that geographical limits are formed by physiological limits on population growth imposed by the environment. And as such, the range limits are determined by bioclimatic conditions for which the population replacement rate is zero (Holt et al. 2005). Based on this logic, in a landscape with a broadscale environmental gradient, the species range limit is aligned with the bioclimatic limit (von Humboldt and Bonpland 1807). Hence, the bioclimatic theory of species distribution is, in essence, a framework on how to project Hutchinson’s (1957) fundamental niche onto the geographical space (Pearson and Dawson 2003).

However, it has long been recognized that the match between the realized environment and fundamental niche may be imperfect. For instance, dispersal from a high-quality source habitat can maintain the population in a sink that would otherwise go locally extinct (Pulliam 1988). Therefore, source-sink dynamics can generate a mismatch between geographical and bioclimatic limits (Pulliam 2000). These observations stand in sharp contrast to the assertions of bioclimatic modelers who claim that at broad spatial scales, the climate is the primary determinant of species distribution (Pearson and Dawson 2003) and, as such, biogeographers can ignore potential mismatches between geographical and bioclimatic limits due to dispersal (Phillips et al. 2006). The main justification of this idea is that—occupied sink patches must be linked to source patches at a length scale comparable to the mean dispersal distance of an organism (Shmida and Wilson 1985, Tittler et al. 2006). And since mean dispersal distance is typically orders of magnitude smaller than distribution length scales, the mismatch between range and bioclimatic limits due to source-sink dynamics is small and can therefore be neglected (also see Boulangeat et al. 2012).

This argument, however, has a significant limitation because it assumes sink patches do not produce migrants despite several studies indicating otherwise. For example, Kanda et al. (2009) noted that Virginia opossum (*Didelphis virginiana*) in Massachusetts used sink habitats near the bioclimatic limit as stepping-stones to occupy sink patches at a distance much greater than mean dispersal distance, thereby extending the range limit far beyond the bioclimatic envelope. In another study, Goel et al. (2020) showed that stepping-stone dispersal, coupled with bistable dynamics, could explain the climatic deviation of the continental-scale savanna-forest boundary from its bioclimatic limit.

These studies highlight that interlinkage of population dynamics of faraway populations by intermediate patches through dispersal, here referred to as stepping-stone dynamics, for short, could be an important determinant of distribution limits. However, we lack an understanding of how demographic processes—such as birth, death, immigration, and emigration—interact at species boundary to drive the mismatch between the geographical and bioclimatic limits. Furthermore, the lack of a mechanistic model renders us unable to ascertain when and where stepping-stone dynamics may be important. Here we address this gap by considering a two-dimensional reaction-diffusion model that combines environment-dependent growth and dispersal characteristics of the species. We discuss the implications of our findings on relating an organism’s niche to distribution, which conceptually links range limit and bioclimatic theory, and underlies our efforts to predict the distribution patterns and make biogeographic inferences (Colwell and Rangel 2009, Peterson et al. 2011).

## Methods

We consider a species whose growth is limited by climate, such that the per-capita growth rate, *r*, is a scalar function of the local environment, *e*. Although, in reality, most species are limited by multiple environmental variables, to build intuition, we present results for a single limiting environment. In Appendix S1 in Supporting Information, we provide results for a species limited by multiple environmental variables.

We assume an increasing gradient in the environment along the *x* direction on a 2D landscape, such that when *e* < *e*\*, the local growth rate is positive, and when *e* > *e*\*, the local growth rate is negative. Here, *e*\* is the bioclimatic limit of the species. More precisely, in two spatial dimensions, the bioclimatic limit forms a contour corresponding to *r*(*e*\*) = 0 (black contour in Fig. 1), bounding where the species exhibits a positive growth rate. Therefore, in the absence of dispersal, the species is present in source locations where the population is above the replacement rate (*i.e.*, *r* > 0) and is absent in sink locations where the population is below the replacement rate (*i.e.*, *r* < 0).

**Figure 1.**
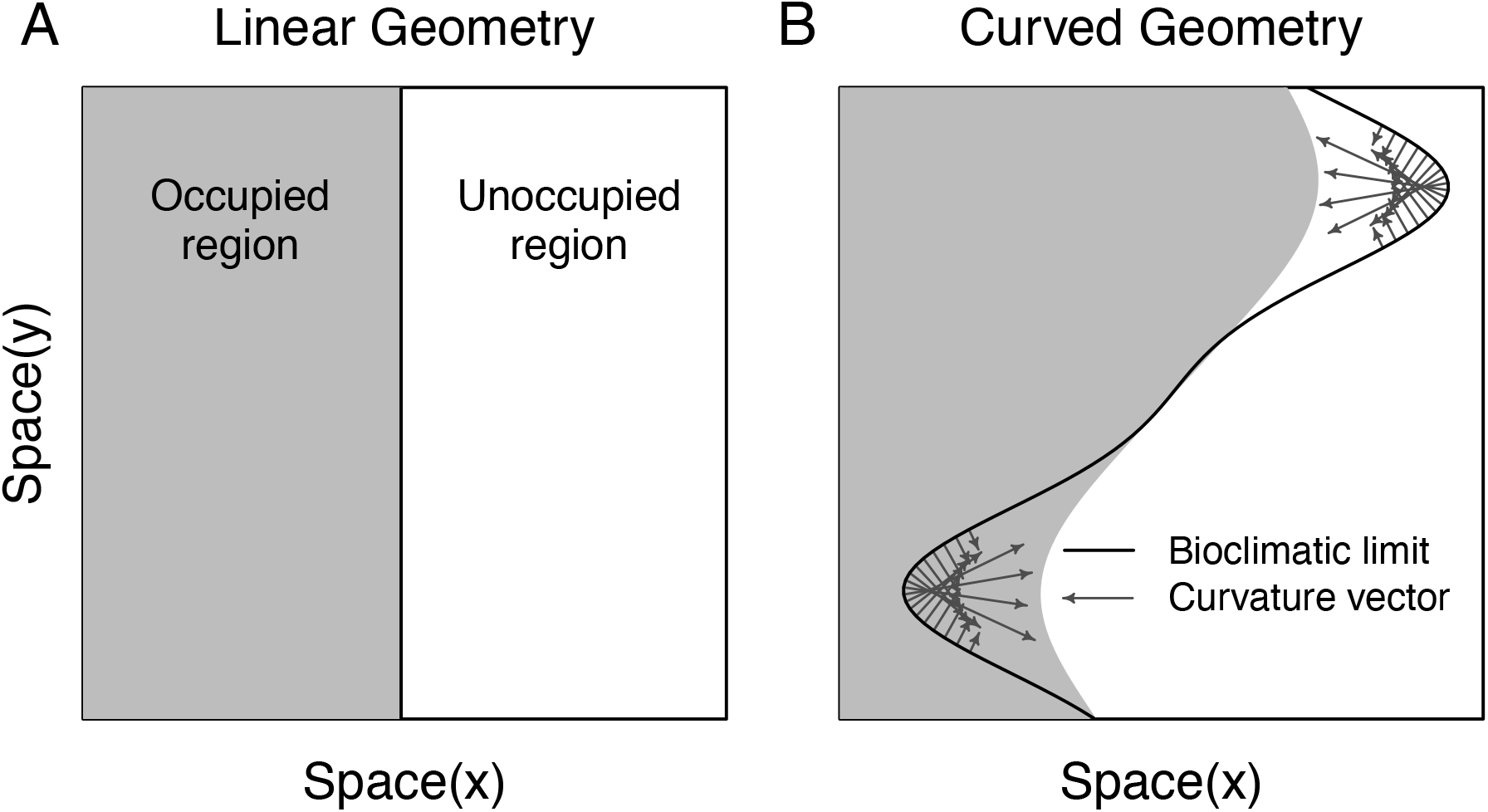
Distribution of species in a landscape with a increasing gradient in environment along *x* direction for (A) linear and (B) curved shapes of the bioclimatic limit (black line). When the bioclimatic limit is a straight line (A), the range limit aligns with the bioclimatic limit because of immigration balances emigration. However, when the bioclimatic limit is curved (B), immigration and emigration rates may differ. As a result, the range limit deviates from the bioclimatic limit in the direction of the curvature vector (grey arrow), with magnitude determined by the growth and dispersal characteristics of the species (see Eq. 3). The length of grey arrows is proportional to the environmental mismatch at the bioclimatic limit and the range limit. The grey region indicates occupied patches. The simulations were performed on a 2D lattice of size 150×150 using Euler forward-time scheme with parameters *∂r*/*∂e*|_*e*=*e*_ = 0.1, *e*\* = 5, *D* = 25, Δ*x* = 1, and Δ*t* = Δ*x*^2^/5*D*.

Next, we incorporate dispersal via the diffusion approach. The diffusion model has an advantage in that it is analytically tractable (Holmes et al. 1994) and emulates observed dispersal patterns among plants and animals (Okubo and Levin 2001). Mathematically, we can express the joint contribution of growth and dispersal as a partial differential equation that captures variation in the population (*N*) in both space and time:

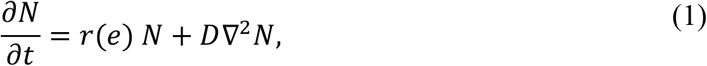

where *D* is the dispersal rate of the species and ∇^2^ (= *∂*^2^/*∂x*^2^ + *∂*^2^/*∂y*^2^) is the diffusion operator that approximates the spatially structured dispersal process in a 2D landscape. Based on this model formalism, the mean dispersal distance (*σ*) of the species is 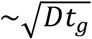, were *t_g_* is the generation time of the species. Although we consider an exponential growth term, our results are robust to non-linearities in growth rate; one can simply replace *r* with *∂f*(*N*, *e*)/*∂N*|_*N*=0_ in the derivations to account for non-linearities, where *f*(*N*, *e*) is the density-dependent growth rate.

## Results

Population change, including at species ecotone, is regulated by four processes: birth and death (captured by the exponential growth term), and immigration and emigration (captured by the diffusion term). Although the growth rate at the bioclimatic limit is zero, the boundary location may still deviate from the bioclimatic limit due to immigration and emigration. The boundary stabilizes where the ecological processes that increase boundary population—birth and immigration—balance the ones that decrease the boundary population—death and emigration.

Using a traveling wave solution (Brazhnik and Tyson 1999) for equation (1), we show the above ecological condition is met when dispersal and growth rates at the range limit are related as

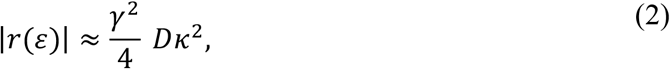

where *ε* is the local environment at the range limit, *γ*^2^ is a positive constant that is independent of species characteristics, and *κ* is the curvature of the bioclimatic limit (see Appendix S1).

When *κ* is small, the mismatch between range and bioclimatic limits in niche space is

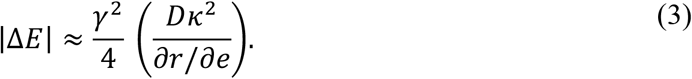

where Δ*E* = *ε* − *e*\* is the difference between the environment at the range and bioclimatic limits, and *∂r*/*∂e* is the sensitivity of species growth rate to changes in the environment at the bioclimatic limit. To interpret these theoretical results, we consider three environmental gradient scenarios. In each scenario, we start with the boundary initially aligned with the bioclimatic limit. Thus, any deviation of the boundary from the bioclimatic limit is solely due to source-sink dynamics.

First, we consider a linear geometry of the bioclimatic limit (*i.e.*, *κ* = 0, Fig. 1A). For this geometry, the number of sink and source patches on either side of the species boundary are equal. As a result, immigration and emigration rates cancel each other to yield net-zero dispersal flux at the boundary. In this trivial case, the range limit aligns with the bioclimatic limit (*i.e.*, Δ*E* = 0, Fig. 1A), which is consistent with our theoretical prediction in equation (3).

However, in reality, the bioclimatic limit rarely assumes a linear geometry (*i.e.*, *κ* ≠ 0). For instance, when the bioclimatic limit is bent with the convex side facing source habitats (bottom half of Fig. 1B), immigration exceeds emigration due to a relatively higher proportion of sources than sinks. Consequently, the boundary population increases and the boundary transgresses slightly into the sink region (grey arrows in Fig. 1B show the direction of movement). Because the shape of the range limit is still convex after a slight shift in the boundary position, immigration at the boundary from these newly occupied sinks still exceeds emigration. As a result, the boundary will continue to encroach sink habitats due to net positive dispersal flux.

So, how far will the species boundary move before it comes to a halt? That depends on the growth and dispersal characteristics of the species. For the convex shape bioclimatic limit, the population in sinks near the bioclimatic limit increases due to dispersal influx from sources and decreases due to inferior patch quality. If the dispersal rate *D* is high, and the quality of sinks declines slowly as one moves away from the bioclimatic limit (*i. e*., *∂r*/*∂e* is low), the sink populations will increase rapidly. As a result, sinks near the bioclimatic limit become an exporter of individuals to adjacent patches. Eventually, the boundary will equilibrate when the positive dispersal flux is balanced by a decrease in growth due to declining patch quality. In this way, the species can use sink patches near the bioclimatic limit as stepping stones to occupy neighboring sinks, thereby extending the distribution far beyond the bioclimatic limit (*i.e.,* Δ*E* ≠ 0; see Eq. 3).

For the third scenario, we consider a bioclimatic limit with its concave side facing source habitats (upper half of Fig. 2B). In contrast to the previous scenario, emigration exceeds immigration and, as a result, the boundary moves backward into source habitats (grey arrows in the upper half of Fig. 2B). Here, too, the magnitude of mismatch between range and bioclimatic limits is determined by dispersal and growth. If the quality of source patches increases slowly and the dispersal outflux is high, the source populations near the bioclimatic limit decrease rapidly to local extinction. This local extinction event creates a domino effect, wherein the boundary continues to encroach source patches until the negative dispersal flux is balanced by an increase in growth rate. Although this scenario may seem counterintuitive as the species range contracts even though source patches are accessible to the species, local extinction due to curved geometry is widely studied in range limit theory (see critical patch size in Okubo and Levin 2001).

Next, we synthesize our results by partitioning the relative contribution of climate and dispersal in determining distribution limits in niche space. Dividing equation (3) by *e*\* we get

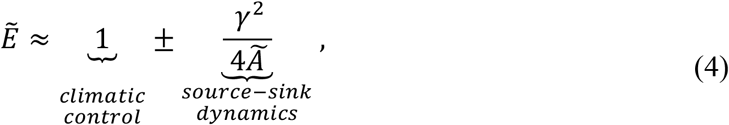

where 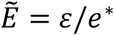 is the rescaled environment at the range limit and

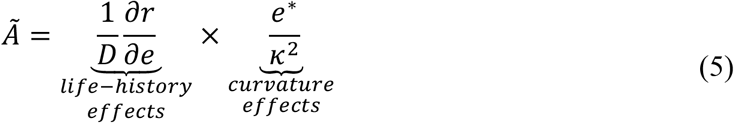

is defined as a dimensionless area that depends on species growth and dispersal and on the curvature of the bioclimatic limit. Rescaling the environment at the range limit offers two important insights. First, we can partition the environment at a range limit for any species into niche constraints imposed by the environment and local source-sink dynamics (see Eq. 4). Second, we can interpret *Ã* as a measure of scale to infer the relative importance of environment and dispersal. When *Ã* is large, the local environment at range limit approaches the bioclimatic limit 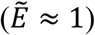. However, when *Ã* is small, the environment at the range limit deviates substantially from the bioclimatic limit.

Although in our derivations, we show the mismatch between range and bioclimatic limits in niche space, we can also express the mismatch in the geographical space. For instance, consider a landscape with an environmental gradient, *G*, in the radial direction. Using equation (3), we can show the mismatch between range and bioclimatic limits in the geographical space is

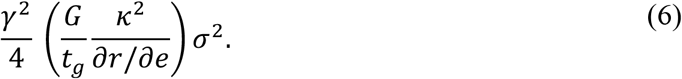

Equation (6) suggests that depending on the species life-history traits, the shape of the bioclimatic limit, and the environmental gradient, the local mismatch in geographical space can be much greater than the species’ mean dispersal distance, *σ*, as illustrated in Fig. 1.

## Discussion

We develop a reaction-diffusion model that mechanistically combines dispersal and growth to examine the role of stepping-stone dynamics in determining distribution limits. We find that species distribution is not only determined by local environmental conditions (niche requirements) but also by source-sink dynamics, which are mediated by the geometrical shape of the bioclimatic limit (Fig. 1 and Eq. 3). For species with high dispersal rate and low growth sensitivity, the mismatch between geographical and bioclimatic limits may be substantial even at large spatial scales (Eq. 6). Based on our analytical results, we propose a dimensionless area, *Ã*, that can be used as a measure of scale to infer the relative importance of dispersal and climate in determining range limits (Eq. 4). Our findings raise conceptual and practical challenges for using bioclimatic models in predicting the distribution of biota and generating biological insights.

Conceptually, bioclimatic models work in two steps. First, the species’ fundamental niche, or the bioclimatic envelope, is estimated either by biophysical experiments (Crozier and Dwyer 2006) or by a correlative approach that map species occurrence to prevailing climatic conditions (Phillips et al. 2006). Next, the constructed niche is transferred onto geography, either at a different time or space (Randin et al. 2006). The predicted distribution patterns are then used to make a wide range of biogeographic inferences, such as niche conservatism of invasive species (Liu et al. 2020) and sister taxa (Peterson et al. 1999), range shifts due to climate change (Araujo and Rahbek 2006), finding suitable habitats to introduce endangered species (Martinez-Meyer et al. 2006), discovering new species (Raxworthy et al. 2003) or populations (Feria and Peterson 2002) in unsampled regions, identifying historical refugia (Waltari et al. 2007), mapping risk potential of disease vectors (Moffett et al. 2007), and many more. Naturally, the robustness of these inferences depends on how reliably we can construct niches and reproject them onto geographical space.

The diffusion model suggests that, for certain life histories, the geographical limit may deviate substantially from the bioclimatic limit due to stepping-stone dynamics (see Fig. 1). As a result, ignoring dispersal can lead to systematic errors in constructing and transferring species bioclimatic envelope (Gilroy and Edwards 2017) even at large spatial scales. First, when estimating bioclimatic envelope using occurrence data, the model may include the environment from occupied sinks and fail to capture the environment in empty sources. Second, even if the bioclimatic envelope is known (e.g., via biophysical experiments), transferring the envelope to a different region may lead to projection errors near the bioclimatic limit as the model may fail to predict occupied sinks and empty sources that are accessible through local dispersal. These errors arising from the interactions between growth and dispersal at the population margins can thwart forecasting efforts and yield misleading inferences.

Although the evidence for large-scale source-sink dynamics is scant, in a recent study, Goel et al. (2020) showed that stepping-stone dispersal at the savanna-forest boundary could explain the continental-scale mismatch between biome and bioclimatic limits. In particular, the authors found that the mismatch in local precipitation at the biome boundary in Africa and the bioclimatic limit was consistent with curvature dynamics predicted from theory. However, these results should be interpreted with caution as curvature dynamics have not been replicated on other tropical continents, and it is always possible that latent variables could produce observed biome patterns. Nevertheless, source-sink dynamics at the savanna-forest boundary in Africa raise two points. First, source-sink dynamics may operate at length scales 10: to 10; times greater than typically assumed (Shmida and Wilson 1985). Second, ignoring stepping-stone dispersal at bioclimatic limit may bias biogeographic inferences about the mechanisms that maintain geographic limits.

It is, therefore, prudent to ask—is bioclimatic theory useful? We think it is. After all, there is overwhelming evidence that the distribution of biota, both in the past (Davis and Shaw 2001) and present (Walther et al. 2002), closely tracks climate, albeit the match is not always perfect (Araujo and Peterson 2012). Instead, we advocate for the cautious use of bioclimatic theory to project distribution patterns and make biological inferences. Our analysis (Eq. 3) suggests that bioclimatic models may be suitable for species sensitive to changes in the environment and have a low dispersal rate. In contrast, for species with fast dispersal and low growth sensitivity, stepping-stone dynamics may be substantial and, therefore, a correlative bioclimatic model may yield erroneous inferences—even at large spatial scales. For such species, one way forward would be to develop Dynamic Range Models that statistically integrate both niche requirements and dispersal based on observational data (see Schurr et al. 2012).

## Acknowledgments

NG thank Vishwesha Guttal for his mentorship and pedagogical training, Lokesh Mishra for discussion on the use of dimensional analysis, Stephen C. Stearns for his commitment to teaching writing, and Abhishek Bhattacharjee, Alvaro Sanchez, and Erika Edwards for their commitment to improving graduate student mentorship. We thank Farrior and Wolf lab members and Eva Arroyo for feedback on the manuscript.

## References

Araujo, M. B., and A. T. Peterson. 2012. Uses and misuses of bioclimatic envelope modeling. Ecology 93:1527–1539.

Araujo, M. B., and C. Rahbek. 2006. How does climate change affect biodiversity? Science 313:1396–1397.

Boulangeat, I., D. Gravel, and W. Thuiller. 2012. Accounting for dispersal and biotic interactions to disentangle the drivers of species distributions and their abundances. Ecology Letters 15:584–593.

Brazhnik, P. K., and J. J. Tyson. 1999. Velocity-curvature dependence for chemical waves in the Belousov-Zhabotinsky reaction: Theoretical explanation of experimental observations. Physical Review E 59:3920–3925.

Colwell, R. K., and T. F. Rangel. 2009. Hutchinson’s duality: the once and future niche. Proceedings of the National Academy of Sciences 106:19651–19658.

Crozier, L., and G. Dwyer. 2006. Combining population-dynamic and ecophysiological models to predict climate-induced insect range shifts. The American Naturalist 167:853–866.

Davis, M. B., and R. G. Shaw. 2001. Range shifts and adaptive responses to Quaternary climate change. Science 292:673–679.

Feria, T., and A. Peterson. 2002. Using point occurrence data and inferential algorithms to predict local communities of birds. Diversity and distributions 8:49—56.

Gaston, K. J. 2003. The structure and dynamics of geographic ranges. Oxford University Press, Oxford.

Gilroy, J. J., and D. P. Edwards. 2017. Source-sink dynamics: a neglected problem for landscape-scale biodiversity conservation in the tropics. Current Landscape Ecology Reports 2:51—60.

Goel, N., V. Guttal, S. A. Levin, and A. C. Staver. 2020. Dispersal increases the resilience of tropical savanna and forest distributions. The American Naturalist 195:833–850.

Holmes, E. E., M. A. Lewis, J. E. Banks, and R. R. Veit. 1994. Partial-Differential Equations in Ecology - Spatial Interactions and Population-Dynamics. Ecology 75:17–29.

Holt, R. D., and T. H. Keitt. 2005. Species’ borders: a unifying theme in ecology. Oikos 108:3–6.

Holt, R. D., T. H. Keitt, M. A. Lewis, B. A. Maurer, and M. L. Taper. 2005. Theoretical models of species’ borders: single species approaches. Oikos 108:18—27.

Kanda, L. L., T. K. Fuller, P. R. Sievert, and R. L. Kellogg. 2009. Seasonal source–sink dynamics at the edge of a species’ range. Ecology 90:1574–1585.

Liu, C., C. Wolter, W. Xian, and J. M. Jeschke. 2020. Most invasive species largely conserve their climatic niche. Proceedings of the National Academy of Sciences 117:23643—23651.

Martinez-Meyer, E., A. T. Peterson, J. I. Servin, and L. F. Kiff. 2006. Ecological niche modelling and prioritizing areas for species reintroductions. Oryx 40:411—418.

Moffett, A., N. Shackelford, and S. Sarkar. 2007. Malaria in Africa: vector species’ niche models and relative risk maps. PloS one 2:e824.

Okubo, A., and S. A. Levin. 2001. Diffusion and ecological problems: modern perspectives. Springer, Berlin, Germany.

Pearson, R. G., and T. P. Dawson. 2003. Predicting the impacts of climate change on the distribution of species: are bioclimate envelope models useful? Global Ecology and Biogeography 12:361–371.

Peterson, A. T., J. Soberon, R. G. Pearson, R. P. Anderson, E. Martinez-Meyer, M. Nakamura, and M. B. Araujo. 2011. Ecological niches and geographic distributions. Princeton University Press, Princeton, New Jersey, USA.

Peterson, A. T., J. Soberon, and V. Sanchez-Cordero. 1999. Conservatism of ecological niches in evolutionary time. Science 285:1265–1267.

Phillips, S. J., R. P. Anderson, and R. E. Schapire. 2006. Maximum entropy modeling of species geographic distributions. Ecological Modelling 190:231–259.

Pulliam, H. R. 1988. Sources, Sinks, and Population Regulation. The American Naturalist 132:652–661.

Pulliam, H. R. 2000. On the relationship between niche and distribution. Ecology Letters 3:349–361.

Randin, C. F., T. Dirnbock, S. Dullinger, N. E. Zimmermann, M. Zappa, and A. Guisan. 2006. Are niche-based species distribution models transferable in space? Journal of Biogeography 33:1689—1703.

Raxworthy, C. J., E. Martinez-Meyer, N. Horning, R. A. Nussbaum, G. E. Schneider, M. A. Ortega-Huerta, and A. T. Peterson. 2003. Predicting distribution of known and unknown reptile species in Madagascar. Nature 426:837—841.

Schurr, F. M., J. Pagel, J. S. Cabral, J. Groeneveld, O. Bykova, R. B. O’Hara, F. Hartig, W. D. Kissling, H. P. Linder, G. F. Midgley, B. Schröder, A. Singer, and N. E. Zimmermann. 2012. How to understand species’ niches and range dynamics: a demographic research agenda for biogeography. Journal of Biogeography 39:2146–2162.

Shmida, A., and M. V. Wilson. 1985. Biological determinants of species diversity. Journal of Biogeography 12:1–20.

Tittler, R., L. Fahrig, and M.-A. Villard. 2006. Evidence of large-scale source-sink dynamics and long-distance dispersal among Wood Thrush populations. Ecology 87:3029–3036.

von Humboldt, A., and A. Bonpland. 1807. Essai sur la géographie des plantes. Levrault, Schoell et Compagnie, Paris.

Waltari, E., R. J. Hijmans, A. T. Peterson, A. S. Nyari, S. L. Perkins, and R. P. Guralnick. 2007. Locating pleistocene refugia: comparing phylogeographic and ecological niche model predictions. PloS one 2:e563.

Walther, G.-R., E. Post, P. Convey, A. Menzel, C. Parmesan, T. J. Beebee, J.-M. Fromentin, O. Hoegh-Guldberg, and F. Bairlein. 2002. Ecological responses to recent climate change. Nature 416:389—395.

