## Appendix S1 for "Source-sink dynamics can maintain mismatched range and bioclimatic limits even at large spatial scales"

**Appendix S1: Calculations and Derivation**

Brazhnik and Tyson (1999) showed that in a homogenous landscape with fixed $r$ and $D$, reaction-diffusion model in equation (1) has a traveling-wave solution, where the boundary speed, $v$, is given by the following relation:

$$v\left[1-\left(\frac{v}{v_0}\right)^2\right] = 2\gamma D\kappa. \tag{A1}$$

Here, $v_0 = 2\sqrt{|r|D}$, $\gamma$ is a positive constant, $\kappa$ is the local curvature of the boundary, and $D$ is the dispersal constant. However, in main text, we assume an inhomogeneous landscape with gradient in $r$. To resolve this issue, we make an additional assumption that gradient in environment is shallow. Therefore, the speed of the boundary can be assumed to be in quasi-equilibrium, such that the instantaneous $v$ is still given by equation (A1). Next, assuming small $\kappa$, we simplify equation (A1) to $v \approx v_o - \gamma D\kappa$. Next, to find environmental conditions at the range limit, we set $v = 0$, which yields $v_o \approx \gamma D\kappa$. Plugging the expression for $v_o$, we find that the per-capita growth rate at the range limit is

$$r \approx \pm\frac{\gamma^2}{4} D\kappa^2. \tag{A2}$$

Next, we Taylor expand growth rate, $r$, around the bioclimatic limit, $\boldsymbol{e}^* = (e_1^*, e_2^* \dots, e_n^*)$, to find the environmental conditions at the range limit, $\boldsymbol{\varepsilon} = (\varepsilon_1, \varepsilon_2, \dots \varepsilon_n)$:

$$\boldsymbol{\nabla} r. \Delta\boldsymbol{E} \approx \pm\frac{\gamma^2}{4} D\kappa^2, \tag{A3}$$

where

$$\Delta\boldsymbol{E} = \sum_k (\varepsilon_k - e_k^*)\widehat{\boldsymbol{e_k}}$$

is the position vector corresponding to the range limit and

$$\boldsymbol{\nabla} r = \sum_k \frac{\partial r}{\partial e_k}\bigg|_{\boldsymbol{e}=\boldsymbol{e}^*} \widehat{\boldsymbol{e_k}}$$

is the gradient vector of $r$ evaluated at the bioclimatic limit. Here $\widehat{\boldsymbol{e_k}}$ is the $k^{th}$ unit environmental vectors in the $n$-dimensional niche space. Finally, we solve equation (A3) to get

$$\Delta\boldsymbol{E} \approx \pm\frac{\gamma^2}{4}\left(\frac{D\kappa^2}{||\boldsymbol{\nabla} r||}\right)\left(\frac{\boldsymbol{\nabla} r}{||\boldsymbol{\nabla} r||}\right), \tag{A4}$$

where $||\quad||$ is the Euclidian norm and $\boldsymbol{\nabla} r/||\boldsymbol{\nabla} r||$ is the normalized gradient vector of $r$. Note, when $n = 1$, we recover equation (3) in the main text. More importantly, our analysis reveals that the environment at the range limit deviates from the bioclimatic limit in the direction of the steepest change in $r$ with magnitude that depends on the dispersal ($D$) and growth ($\|\nabla r\|$) characteristic of the species and on the geometrical shape ($\kappa^2$) of the bioclimatic limit.

Note, here we use $\kappa$ as the curvature of both range and bioclimatic limits. Although, strictly, $\kappa$ is the curvature of the range limit but can be approximated as the curvature of the bioclimatic limit. See in Fig. 1, the range limit bends in the same direction as the bioclimatic limit, with magnitude either equal (Fig. 1A) or slightly less (Fig. 1B).

**References**

Brazhnik, P. K., and J. J. Tyson. 1999. Velocity-curvature dependence for chemical waves in the Belousov-Zhabotinsky reaction: Theoretical explanation of experimental observations. Physical Review E **59**:3920-3925.
